# Class I DISARM provides anti-phage and anti-conjugation activity by unmethylated DNA recognition

**DOI:** 10.1101/2021.12.28.474362

**Authors:** Cristian Aparicio-Maldonado, Gal Ofir, Andrea Salini, Rotem Sorek, Franklin L. Nobrega, Stan J.J. Brouns

## Abstract

Bacteriophages impose a strong evolutionary pressure on microbes for the development of mechanisms of survival. Multiple new mechanisms of innate defense have been described recently, with the molecular mechanism of most of them remaining uncharacterized. Here, we show that a Class 1 DISARM (defense island system associated with restriction-modification) system from *Serratia* sp. provides broad protection from double-stranded DNA phages, and drives a population of single-stranded phages to extinction. We identify that protection is not abolished by deletion of individual DISARM genes and that the absence of methylase genes *drmMI* and *drmMII* does not result in autoimmunity. In addition to antiphage activity we also observe that DISARM limits conjugation, and this activity is linked to the number of methylase cognate sites in the plasmid. Overall, we show that Class 1 DISARM provides robust anti-phage and anti-plasmid protection mediated primarily by *drmA* and *drmB*, which provide resistance to invading nucleic acids using a mechanism enhanced by the recognition of unmethylated cognate sites of the two methylases *drmMI* and *drmMII*.

## INTRODUCTION

The arms race between prokaryotes and bacteriophages drives their co-evolutionary dynamics and has led to the evolution of multiple antiviral defense mechanisms in prokaryotes that are collectively known as the prokaryotic immune system (1). Most prokaryotic defenses are innate, acting via recognition of general signals that are not shared by the microbial cell. A well-known example of innate defense are the highly abundant restriction-modification (R-M) systems (2) that cleave phage nucleic acids at sequence motifs protected in the host chromosome by epigenetic modifications (3). Recently, the analysis of genomic neighborhoods of known defense systems revealed multiple new anti-phage systems (4–12), among which the defense island system associated with restriction-modification (DISARM) (13).

The DISARM system is composed of three-core genes: gene *drmA* with a helicase domain (pfam00271); gene *drmB* with a DUF1998 domain (pfam09369, helicase-associated); and gene *drmC* with a phospholipase D (PLD) domain (pfam13091). In the uncharacterized Class 1 DISARM systems, this core triplet is preceded by the SNF2-like helicase *drmD* (pfam00176), and the DNA adenine N6 methylase *drmMI* (pfam13659) (13). Class 2 DISARM systems contain, in addition to the core gene triplet, the DNA 5-cytosine methylase *drmMII* (pfam00145) and, on occasion, the gene of unknown function *drmE.* The Class 2 DISARM system was shown to use methylation of specific host motifs to mark self-DNA, akin to R-M systems, but its molecular mechanism remains largely unknown. Unlike R-M systems, DISARM seems to not fully depend on the sequence motif identified by the methylase to interfere with the incoming DNA; importantly, the candidate nuclease of the system (*drmC*) was found dispensable for resistance and its activity is yet unknown.

In some cases, Class 1 DISARM systems are accompanied by an additional cytosine methylase *drmMII* gene (13). Here we characterized such a Class 1 DISARM system present in *Serratia* sp. SCBI (**Figure 1A**). We found that Class 1 DISARM provides broad anti-phage and anti-conjugation activity independent of methylation status of the incoming DNA, and drives a population of chronic infecting phages to extinction. Unlike Class 2 DISARM and R-M systems, the methylases of Class 1 DISARM only partially methylate adenine and cytosine bases of the host DNA at motifs ACAC<u>A</u>G and MT<u>C</u>GAK, and the absence of the methylases does not result in autoimmunity. Overall, our results show that DISARM combines methylation and non-methylation signals to provide protection against invader DNA, establishing a clear distinction from R-M systems.

**Figure 1.**
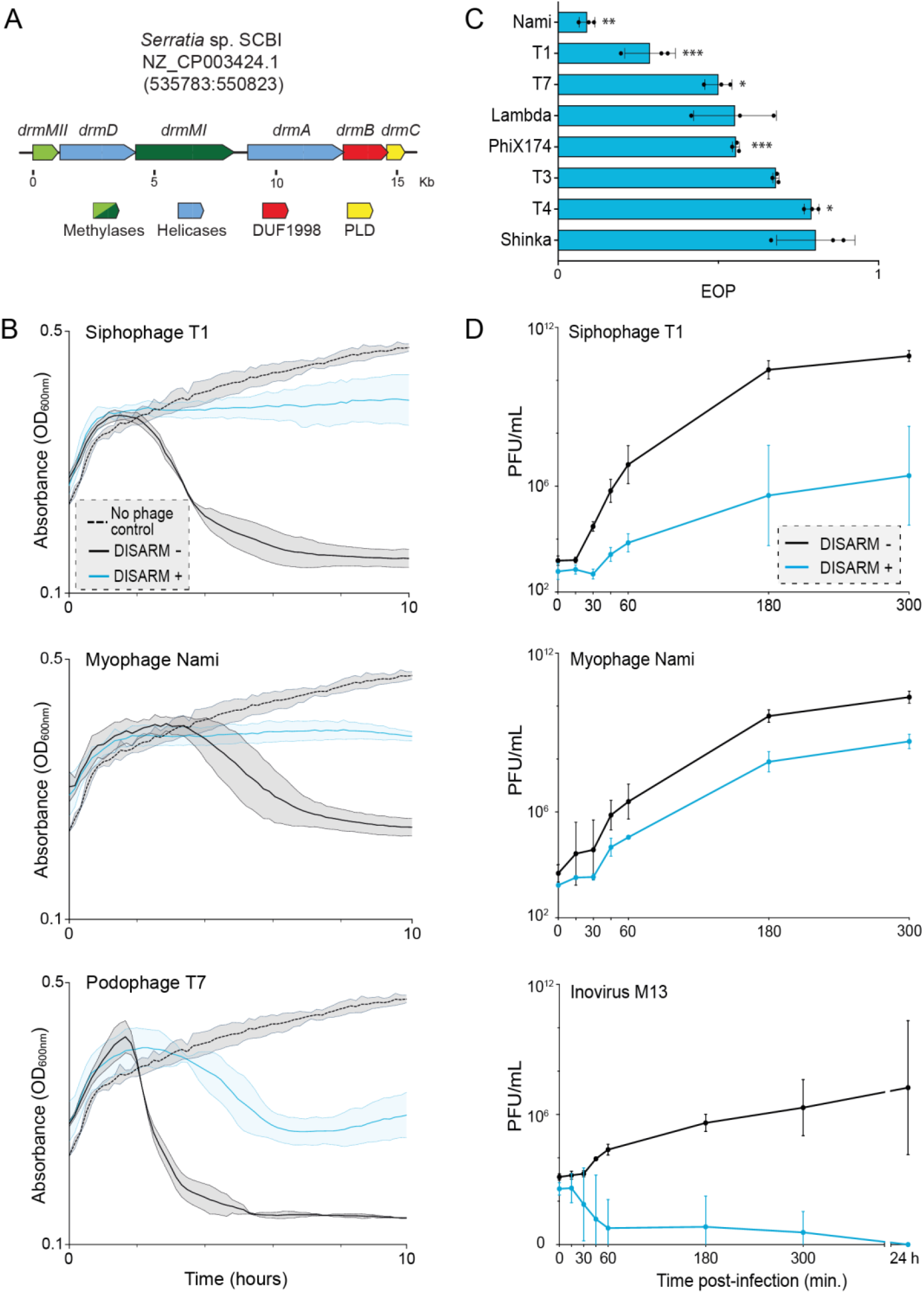
Protection provided by Class 1 DISARM against phage infection. (**A**) Gene cluster of Class 1 DISARM from *Serratia* sp. SCBI. (**B**) Effect of siphophage T1, myophage Nami, and podophage T7 on the growth curve of DISARM (+) or DISARM (-) strains. Uninfected DISARM (-) and DISARM (+) strains have similar growth curves and only uninfected DISARM (-) is displayed. Full results can be seen in **Supplementary Figure S2**. Initial MOI: 5×10^-4^. Filled areas inside dotted lines indicate standard deviation of three independent replicates. (**C**) Efficiency of plating (EOP) of a set of DNA phages in a DISARM (+) strain normalized to the DISARM (-) strain. (**D**) Effect of DISARM on the population of siphophage T1, myophage Nami, and inovirus M13 over time. Bacterial cultures of DISARM (-) or DISARM (+) strains were infected with phage at MOI of 2×10^-6^, 1×10^-5^ and 2×10^-6^ for phages T1, Nami and M13 respectively, and the titer was determined at selected time points. Curves depict the average and standard deviation of three independent experiments. Statistical significance was determined by two-tailed, unpaired t Test and is represented as *, **, or *** for p < 0.05, 0.01 or 0.001, respectively.

## MATERIAL AND METHODS

### Bacterial strains and growth conditions

*Escherichia coli* strains DH5α, BL21-AI, JM109, ER2738, WG5, C3000, and S17-1 were cultured in Lysogeny Broth (LB) media at 37°C. For solid media experiments, LB was supplemented with 1.5% (w/v) agar (LBA) and the cultures incubated overnight at 37°C. When required, media was supplemented with antibiotics at the following final concentrations: 100 μg/mL ampicillin, 50 μg/mL kanamycin, 50 μg/mL streptomycin, 50 μg/mL gentamicin, and 25 μg/mL chloramphenicol. Gene expression was induced with 1 mM Isopropyl β-d-1-thiogalactopyranoside (IPTG) and 0.2% (w/v) L-arabinose. The *Serratia* sp. SCBI strain (South African *Caenorhabditis briggsae* Isolate) was grown aerobically in LB media at 30°C with shaking at 180 rpm.

### Phage cultivation

*E. coli* phages were propagated using their host strain (**Supplementary Table S1**) as described previously (14). Briefly, bacterial cultures at early exponential growth phase (0.3-0.4 OD600) were infected with a phage lysate and incubated overnight at 37°C with shaking. Cultures were spun down and the supernatant filtered (0.2 μm PES) and stored at 4°C until use. When required, phages were concentrated by adding PEG-8000 and NaCl at final concentrations of 100 mg/mL and 1 M, respectively. The suspension was incubated overnight at 4°C, centrifuged at 11,000 × *g* at 4°C for 1h, and the phage-containing pellet was re-suspended in SM buffer (100 mM NaCl, 8 mM MgSO_4_.7H_2_O, and 50 mM Tris-HCl pH 7.5). The phage titer was determined using the small drop plaque assay method as described by Mazzocco and colleagues (15). *E. coli* strain BL21-AI was used for titering and performing assays of phages T1, T3, T4, T7, Lambda-vir, Myo21S, and Myo22L, strain WG5 was used for MS2, and strain C3000 was used for PhiX174. Phage M13 was titered using *E. coli* strain ER2738 and 0.6% LBA supplemented with 1 mM IPTG and 200 μg/mL X-gal. Assays of M13 phage were performed in strain JM109 (DE3).

### Cloning of Class 1 DISARM and mutants

Plasmids and primers used in this study are listed in **Supplementary Table S2** and **Supplementary Table S3**, respectively. The complete Class 1 DISARM system and different combinations of its genes were amplified by PCR from *Serratia* sp. SCBI genomic DNA with primers indicated in **Supplementary Table S3** and using Q5 DNA Polymerase (New England Biolabs) according to the manufacturer’s instructions. The PCR products were cloned into the plasmid backbones using the NEBuilder HiFi DNA Assembly Cloning Kit (New England Biolabs) following manufacturer’s instructions. Plasmid pDIS_3 was used as a template for PCR with primers listed in **Supplementary Table S3** to construct pDIS_5, pDIS_6 and pDIS_7 by restriction cloning. All plasmids were confirmed by Sanger sequencing (Macrogen) and transformed into strains *E. coli* BL21-AI and *E. coli* JM109(DE3) for the assays described below.

### Phage infection growth curves

Bacterial cultures with the DISARM system or empty vector were grown to early exponential phase (OD_600_ of 0.2-0.3), induced with IPTG and L-arabinose, and grown for 90 min at 37 °C and 180 rpm. The cultures were normalized to an OD600 of 0.5 (approximately 1×10^8^ CFU/ml) and 190 μl were dispensed into wells of 96-well microtiter plates. Then, 10 μl of phage suspension were added to the wells at different multiplicity of infection (MOI), and bacterial growth was followed in an EPOCH2 microplate reader with OD600 measurements every 10 min at 37°C with constant shaking.

### Phage replication over time

Bacterial cultures were prepared as above and infected with phages at different MOIs. Cultures were incubated at 37°C with shaking at 180 rpm, and phage titers were measured over time using the small drop plaque assay method using the wild type strain as the host.

### Methylation-sensitive DNA sequencing

Genomic DNA was extracted from the wild-type strain and strains containing the complete DISARM system or its methylases, using the phenol chloroform method as described before (16) with some modifications (17). Briefly, cultures at exponential phase were induced and incubated overnight. Bacterial cells were pelleted, re-suspended in TE buffer, and treated with RNase I and lysozyme at 1 μg/ml for 1h, followed by proteinase K at 50 μg/ml for 1h. DNA was extracted twice with phenol:chloroform (1:1) and precipitated by adding 300 mM sodium acetate pH 5.2 and two volumes of absolute ethanol. After incubation at −20°C for 1h or overnight, the DNA was pelleted by centrifugation at 21,000 × *g*, and the pellet re-suspended in nuclease-free water. The DNA was quantified using Qubit dsDNA HS Assay kit (Invitrogen) and the quality assessed using Nanodrop (Thermo Scientific).

The genomic DNA was sequenced by Pacific Biosciences using Single Molecule Real Time (SMRT) sequencing technology to detect DNA methylation sites (18). DNA libraries were prepared using the SMRTbell® Express Template Prep Kit 2.0 and the Barcoded Overhang Adapter kit according to the manufacturer instructions. Sequencing was performed on a PacBio Sequel platform using Sequel sequencing kit 3.0. The data were analyzed using the Base Modification and Motif Analysis module of the SMRT-Link v7.0.1 software to detect methylation motifs.

Bisulfite library preparation and sequencing was done as previously described (13). Sequencing results were analyzed using Bismark v0.20.1 (Krueger & Andrews 2011 https://doi.org/10.1093/bioinformatics/btr167) to identify methylated cytosines. The sequence neighborhood of methylated cytosines was analyzed using Weblogo (Crooks et al. 2004) to determine the methylation motif.

Raw sequencing data were deposited in Figshare for PacBio (doi.org/10.6084/m9.figshare.17295215) and Bisulfite (doi.org/10.6084/m9.figshare.17295212) sequencing.

### Construction of motif-containing conjugative plasmids

Synthetic constructs (Integrated DNA Technologies) of tetracycline-regulated YFP and a motif-adaptable module (MAM) (**Supplementary Table S4**) were introduced into pSEVA_331 by restriction cloning. Putative DISARM motifs were removed from the backbone sequence of pSEVA_331 by replacing these regions with synthetic constructs (**Supplementary Table S4**). For this, pSEVA_331 and the synthetic constructs were amplified using primers in **Supplementary Table S3** and Q5 DNA Polymerase (New England Biolabs), and cloned by restriction digest using the enzymes indicated in **Supplementary Tables S3** and **S4**, giving rise to plasmid pCONJ. Plasmids pCONJ_1 to pCONJ_8 (**Supplementary Table S2**) were created by cloning synthetic constructs (**Supplementary Table S4**) containing different motif combinations into the MAM regions of pCONJ using restriction digest as above. All plasmid constructs were confirmed by sequencing (Macrogen) and transformed into *E. coli* BL21-AI.

### Conjugation efficiency

The *E. coli* strain S17-1 containing variants of the plasmid pCONJ (**Supplementary Table S2**) was used as the donor strain. Cells were grown to exponential phase and induced when necessary. Approximately 5×10^8^ cells of both donor strain and the DISARM or wild-type strain (recipient) were spun down and re-suspended in 5 ml of fresh LB media. The strains were combined in equal counts in a final concentration of 1×10^8^ CFU/mL. After gently mixing, cells were pelleted at 2,000 × *g* for 10 min at room temperature, and incubated at 37°C for 4 h without shaking. The cell mixture was plated onto LBA plates containing different antibiotics to determine the proportion of recipient cells that acquired the plasmid from the donor strain. Conjugation efficiency was estimated as the ratio of plasmid-acquisition events versus the total number of recipient strain cells.

### Statistical analysis

The average values of three biological replicates were reported in the result and supplementary sections. Unpaired two-tailed t Test and one-way analysis of variance (ANOVA) with Dunnett’s post-hoc multiple comparison test were used to compare the means between groups. Confidence intervals were set at 95% (* = p < 0.05; ** = p < 0.001; *** = p < 0.0001). Statistical analysis was performed using GraphPad Prism version 5.0 for Windows.

## RESULTS

### Class 1 DISARM protects against widely diverse DNA phages

To determine whether the predicted Class 1 DISARM system from *Serratia* sp. SCBI provides protection from phage infection, we transplanted the six genes of the system into *E. coli* BL21-AI (**Figure 1A**). We then challenged the DISARM-containing strain (DISARM (+)) with *Caudovirales* of three morphologies (sipho-, myo-, and podophages) (19) at different MOI. Class 1 DISARM shows clear anti-phage protection against all phages tested, by preventing or delaying the collapse of the bacterial population upon phage infection even at high MOI (**Figure 1B**, **Supplementary Figure S1**). To quantify the level of protection, we measured the efficiency of plating (EOP) of the same set of phages on the DISARM (+) strain in comparison to control (DISARM (-)) cells. Class 1 DISARM provided significant protection against phages T1, T4, T7, Nami and phiX174 (**Figure 1C**).

Overall, both liquid and solid media assays demonstrate the broad anti-phage activity provided by Class 1 DISARM.

### Class 1 DISARM can drive a phage population with chronic lifestyle to extinction

We next investigated the effects of the Class 1 DISARM on the propagation of a phage and accumulation of active phage in the cell culture. For this, we measured the phage titers periodically upon infection of DISARM (+) or DISARM (-) strains with T1 or Nami phages. The number of infectious T1 phages in the population is reduced by approximately 3 orders of magnitude in DISARM containing strains from 30 min on, reaching a maximum reduction of 5.6×10^4^ fold at 180 minutes post-infection (**Figure 1D**). DISARM also inhibits the propagation of phage Nami, with a maximum reduction of approximately 50-fold at 180 minutes post-infection (**Figure 1D**).

Virulent *Caudovirales* follow a lytic life cycle where the production of phages occurs typically within 10-30 minutes after the ejection of the phage genome, ultimately resulting in cell death for the release of the newly formed phage particles. Phages that follow a chronic life cycle are able to produce new phages continuously without causing cell death, with the new virions extruding out of the cell (20). To investigate the effect of Class 1 DISARM on the propagation of a phage population with chronic lifestyle, we monitored the phage titers of a culture infected with the single-stranded DNA (ssDNA) inovirus M13. We observed a rapid decrease of the number of phages, with DISARM containing strains producing no more phage after 24h (**Figure 1D**), which was not observed for any of the *Caudovirales* tested. To understand if the strong activity of Class 1 DISARM against phage M13 is a consequence of its chronic lifestyle or the type of genetic material (ssDNA versus the double-stranded DNA of *Caudovirales*), we additionally tested DISARM against ssDNA phage phiX174. PhiX174 uses a mechanism of phage DNA replication similar to that of phage M13 (21, 22), but follows a lytic life cycle. The protective effect of DISARM against infection by phiX174 was similar to that obtained for the dsDNA phages (**Supplementary Figure S3A**), suggesting that the strong effect of DISARM against phage M13 is not related to the single stranded nature of the phage genome in the phage particle. We further tested the protective effect of Class 1 DISARM against single-stranded RNA (ssRNA) phage MS2 but observed no protection (**Supplementary Figure S3B**), suggesting that Class 1 DISARM is a DNA-directed defense system. In summary, Class 1 DISARM reduces the number of infectious phages produced over time in a bacterial culture, and is able to completely abolish the propagation of chronic phage M13.

### Class 1 DISARM provides protection independent of methylation status

We next studied the essentiality of the individual genes of the Class 1 DISARM system in anti-phage protection. First, we used EOP assays to challenge cells harboring the complete or partial DISARM system. We observed that DISARM provides protection against infection by dsDNA phages T1 and Nami, and that some of the DISARM genes can be deleted while retaining either full or partial protection (**Figure 2A**, **Supplementary Figure S4A**). To better evaluate gene essentiality in Class 1 DISARM, we followed phage propagation over time in liquid culture of cells harboring the complete or partial DISARM system. Results are shown in **Figure 2B,C and Supplementary Figure S4B** as the increase of phage titer over time at the time point where DISARM achieved the strongest effect compared to the titer at the start of infection. In contrast to what was previously observed for Class 2 DISARM systems (13), deletion of core genes *drmABC* did not abolish the full protective effect for both phages T1 and M13, and only the additional deletion of *drmMII* resulted in complete loss of protection with full restoration of the phage replication capacity. For phage Nami, deletion of core *drmABC* or deletion of *drmD* and *drmMI* resulted in almost complete loss of protection (**Figure 2B**), with full restoration of phage replication being achieved with their combined deletion. Importantly, we observed that *drmC* is not required for the anti-phage activity (**Supplementary Figure S5**) of Class 1 DISARM, as previously observed for Class 2 DISARM (13).

**Figure 2.**
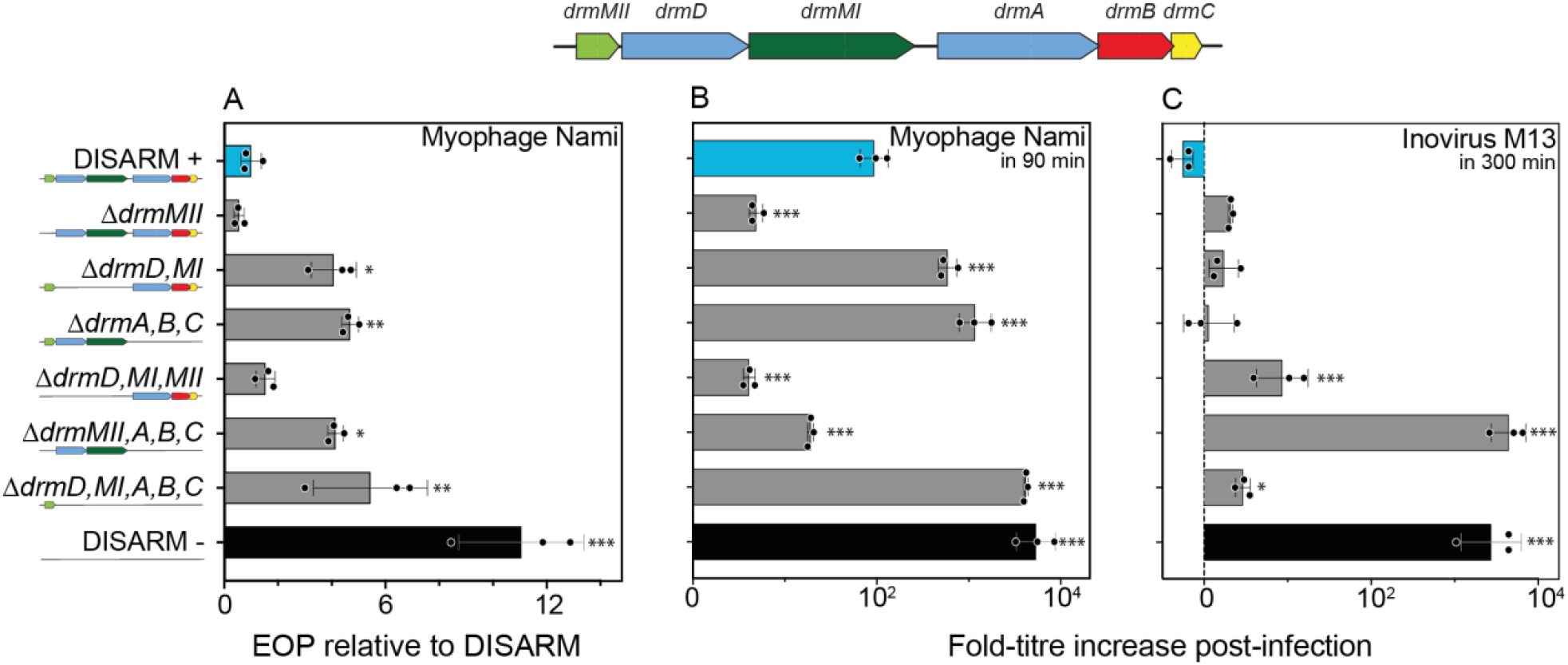
Effect of Class 1 DISARM components on protection against phage infection. (**A**) Efficiency of plating (EOP) of myophage Nami on strains containing the modified DISARM system (DISARM (-) strain), normalized to the DISARM strain. Titer-fold increase of (**B**) myophage Nami, and (**C**) inovirus M13 upon propagation in cultures of strains containing the complete or modified DISARM system. Graphics represent the time point at which maximum effect on phage replication was achieved. For (**C**), negative values indicate phage titers below the initial phage titer. Bars depict the average and standard deviation of three independent replicates. Statistical significance was determined by one-way ANOVA+ with Tukey post-hoc test and is represented as *, **, or *** for p < 0.05, 0.01 or 0.001, respectively.

Overall, we observed that Class 1 DISARM phage protection is relatively robust to deletion of individual genes of the operon.

### Class 1 DISARM of *Serratia* sp. SCBI modifies host DNA with two methylation patterns

To understand if the methylase genes *drmMI* and *drmMII* methylate the host DNA, we sequenced the genomes of the wild type strain, the DISARM (+) strain, and the strain containing one or both methylases using sequencing methods sensitive to epigenetic marks. The adenine methylation by *drmMI* was characterized by PacBio sequencing and revealed an N6-methyladenosine (m6A) modification of the ACAC<u>A</u>G motif (methylated base underlined, **Figure 3A**, **Supplementary Table S5**). The cytosine methylation by *drmMII* was characterized by bisulfite sequencing and revealed a 5-methylcytosine (5mC) modification of MTCGAK motifs (methylated cytosine underlined, **Figure 3B**, **Supplementary Table S5**), which is a distinct motif from the 5mC modification in C<u>C</u>WGG motifs reported for the Class 2 DISARM system of *Bacillus subtilis* (13). The completeness of modification by the methylases was higher in the presence of the full DISARM system (84.3% m6A, and 67.0% 5mC) than in the presence of the two methylases alone (56.5% m6A, and 34.0% 5mC), suggesting some form of synergetic effect by the methylase pair (**Figure 3D**). Curiously, the presence of both *drmMI* and *drmMII* increased the m6A methylation ratio of *drmMI* to 73.9%, but had no effect on 5mC methylation by *drmMII.* No relation was observed between the distance of methylated or unmethylated ACAC<u>A</u>G and MT<u>C</u>GAK motifs, suggesting that their methylation status was independent of their relative location in the genome. In summary, the Class 1 DISARM system of *Serratia* sp. SCBI uses both adenine and cytosine methylation to modify specific motifs in the bacterial chromosome.

**Figure 3.**
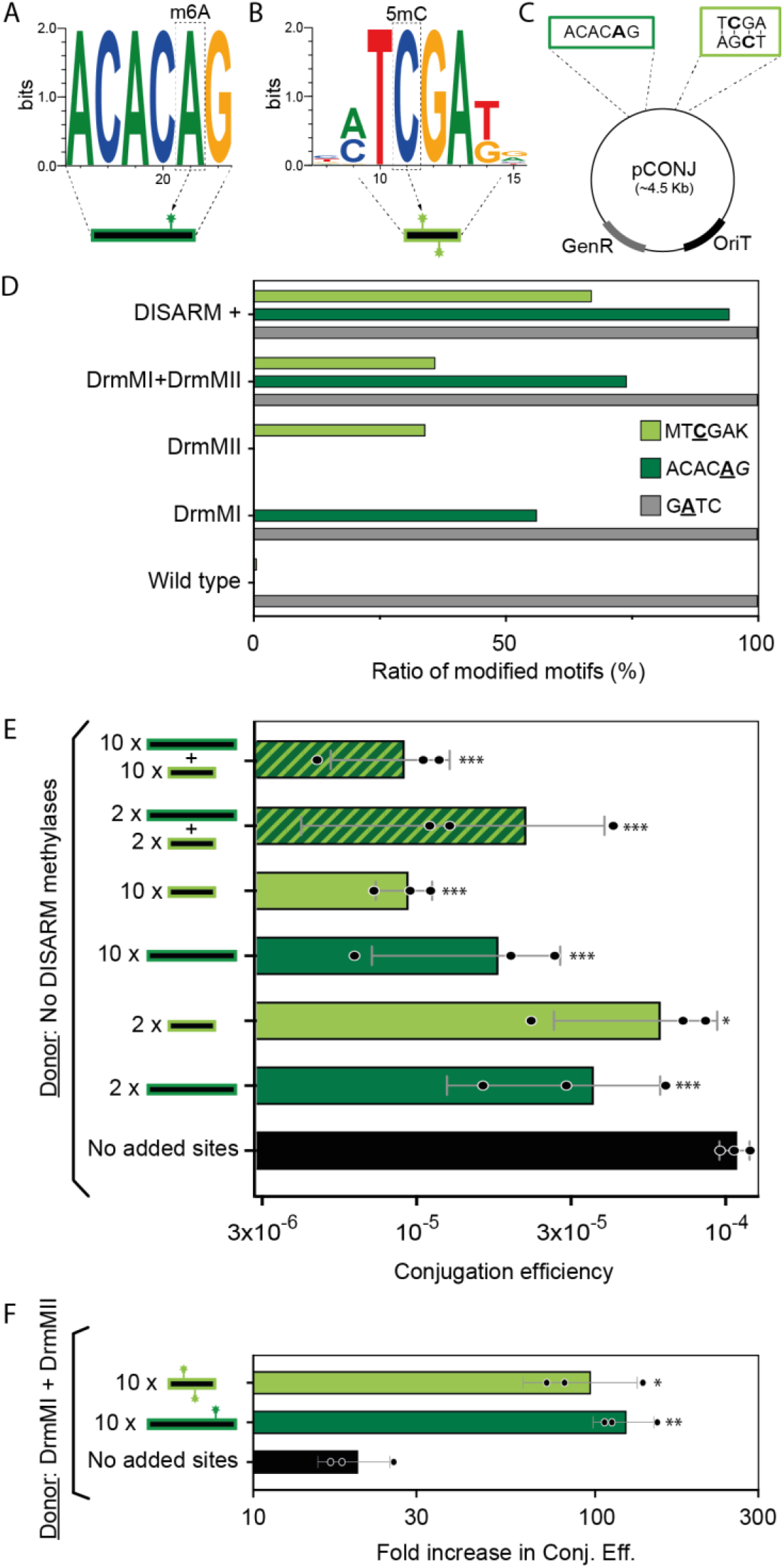
Effect of DNA methylation in Class 1 DISARM. Weblogos of methylation motifs of DISARM methylases (**A**) DrmMI and (**B**) DrmMII. (**C**) Schematic representation of pCONJ plasmid with motifs as defined in (**A**) and (**B**). (**D**) Relative number of modified sites detected in the genome of *E. coli* expressing the Class 1 DISARM system of *Serratia* sp. SCBI. Three distinct DNA modifications were detected: G<u>A</u>TC sites modified by *dam* from *E. coli;* ACACAG sites modified by *drmMI;* and MTCGAK sites modified by *drmMII.* (**E**) Conjugation efficiency of plasmid pCONJ with variable numbers of unmethylated motifs ACAC<u>A</u>G and/or MT<u>C</u>GAK into the recipient DISARM (+) strain (see **Supplementary Figure S5B** for results in recipient control strain). Control is pCONJ with no motifs in its sequence. (**F**) Conjugation efficiency of plasmid pCONJ with 10 methylated motifs ACAC<u>A</u>G or MT<u>C</u>GAK into the recipient DISARM (+) strain, normalized by conjugation efficiency of pCONJ with unmethylated motifs.

### Class 1 DISARM displays anti-conjugation activity dependent on number of cognate sites

To determine the influence of the methylation pattern of the invading DNA on the level of protection by Class 1 DISARM, we performed conjugation assays with plasmid pCONJ which does not contain ACAC<u>A</u>G or MT<u>C</u>GAK motifs (**Supplementary Table S2**). We first assessed the protection provided by DISARM towards conjugation of the unmethylated pCONJ, and observed an approximately 47-fold reduction in conjugation efficiency in the presence of DISARM (**Supplementary Figure S6A**). Next, we engineered different versions of pCONJ with variable numbers of DISARM methylation motifs and performed conjugation assays in the DISARM (+) and DISARM (-) strains as recipients. The conjugation efficiency of non-methylated pCONJ in the DISARM (+) strain decreased moderately with increasing numbers of ACACAG and TCGA motifs present in the plasmid, up to 12-fold (**Figure 3E**, **Supplementary Figure S6B**). The protective effect against plasmid conjugation is only slightly stronger when both types of motifs are present.

Next, we compared the conjugation efficiency of pCONJ originated from cells expressing DISARM methylases DrmMI and DrmMII to that of pCONJ containing non-methylated motifs (**Figure 3F**). The conjugation efficiency of pCONJ was drastically elevated by 134- and 82-fold when the donor strain contained both DrmMI and DrmMII, supporting the role of methylation as an off switch for DISARM activity. The control plasmid, in which no ACACAG and MTCGAK motifs were added, displayed a 21-fold increase in conjugation efficiency with both methylases present in donor strain possibly due to effects of induction of gene expression of both methylases in the donor. Overall, the conjugation assays demonstrate that Class 1 DISARM displays anti-conjugation activity that is enhanced by the presence of unmethylated forms of both methylase motifs in the plasmid.

## DISCUSSION

Here, we show that Class 1 DISARM of *Serratia* sp. SCBI provides broad protection against phages and plasmid conjugation using a mechanism of incoming nucleic acid detection which is enhanced by the recognition of unmethylated cognate sites of the two methylases *drmMI* and *drmMII.* In Class 2 DISARM systems, the deletion of genes *drmA, drmB*, or *drmE* resulted in complete loss of protection (13). The Class 1 DISARM system of *Serratia* sp. provides a more robust protection, in which the deletion of individual genes is not sufficient to abolish the protective effect of the system against some phages (e.g. T1 and M13). We observed that the number of unmethylated motifs present in the incoming foreign DNA increased the protective effect of DISARM against incoming invaders.

We found that the protection against phages was not scaled to the number of *drmMI* and *drmMII* sites in their genomes (**Supplementary Table S6**), suggesting also other factors at play. Similarly, the number of methylation sites on phage genomes did not affect the protection for the BREX (23) system, and the same was suggested for Class 2 DISARM (13). This is markedly distinct from tested R-M systems in which restriction (and therefore protection) is dependent on the number of methylation sites in the invader’s DNA (24). This suggests that DISARM and BREX use mechanisms to identify invader DNA distinct from those of R-M systems, and which may prevent strong negative selection for specific methylation motifs. It is also possible that the intrinsic methylation patterns of phage DNA affect defense by DISARM, as observed previously for R-M and BREX systems (24, 25).

The use of a molecular mechanism distinct from classical R-M systems is further supported by the lack of autoimmunity in cells in the absence of *drmMI* and *drmMII* (**Supplementary Figure S7**), contrary to R-M systems where this results in cleavage of the bacterial DNA (26). This is also consistent with the fact that not all motifs in the bacterial genome of the transplanted host are methylated by the DISARM methylases, unlike the almost complete motif methylation observed in R-M. This suggests a tight regulation of the defense activity of Class 1 DISARM that seems to result from the physical occlusion of the DNA entry site of the DrmAB complex by the trigger loop, which is removed upon presence of a 5’ ssDNA end (27). Interestingly, the methylase drmMII of Class 2 DISARM provides almost complete methylation of motifs in the bacterial chromosome and its absence was found to be deleterious to the cells (13). Differences in the molecular mechanisms of Class 1 and Class 2 DISARM likely result from the use of distinct methylases. We found that *Serratia*’s Class 1 DISARM system has the unique feature of combining methylation of both palindromic and non-palindromic motifs in the bacterial chromosome. Akin to BREX and R-M type I and III systems, DrmMI of Class 1 DISARM methylates a non-palindromic site (ACAC<u>A</u>G). The modification occurs at the adenine in the fifth position of the recognition site, as previously reported for the BREX site TAGGAG (8). Because the recognition motif of DrmMI is non-palindromic, only one DNA strand will be methylated. Some R-M systems (e.g. type III and type ISP) maintain the epigenetic marks by requiring interactions between different sites (28), but it is unclear if DISARM and BREX use similar mechanisms.

Contrary to DrmMI, the DrmMII methylase of Class 1 DISARM modifies a degenerate palindromic site (MT<u>C</u>GAK) in the bacterial chromosome. The methylation site of DrmMII of the *Bacillus paralicheniformis* Class 2 DISARM system was also shown to be palindromic, although of an unrelated sequence (C<u>C</u>NGG) (13), much like the methylases of R-M type II systems. Interestingly, DrmMII (palindromic motif) alone was shown to provide anti-phage activity against the chronic infecting phage M13 and phage T1, possibly by interfering with the phage replication cycle as observed previously (29). DrmMI (non-palindromic motif) had no observable impact on anti-M13 activity, as previously observed also for the BREX system (non-palindromic motif). DrmABC without any of the methylases also provides protection against M13. It is possible that the strong activity of DISARM against phage M13 results from the added effect of DrmABC and a possible direct effect of DrmMII methylation of palindromic sites on DNA replication of M13. In conclusion, we show that Class 1 DISARM systems are effective on viral and plasmid DNA, and show enhanced protection against invader DNA when unmethylated cognate DNA motifs are present (**Figure 4**). The mechanism of DISARM is remarkably robust with many of its components playing an enhancing but non-essential role.

**Figure 4.**
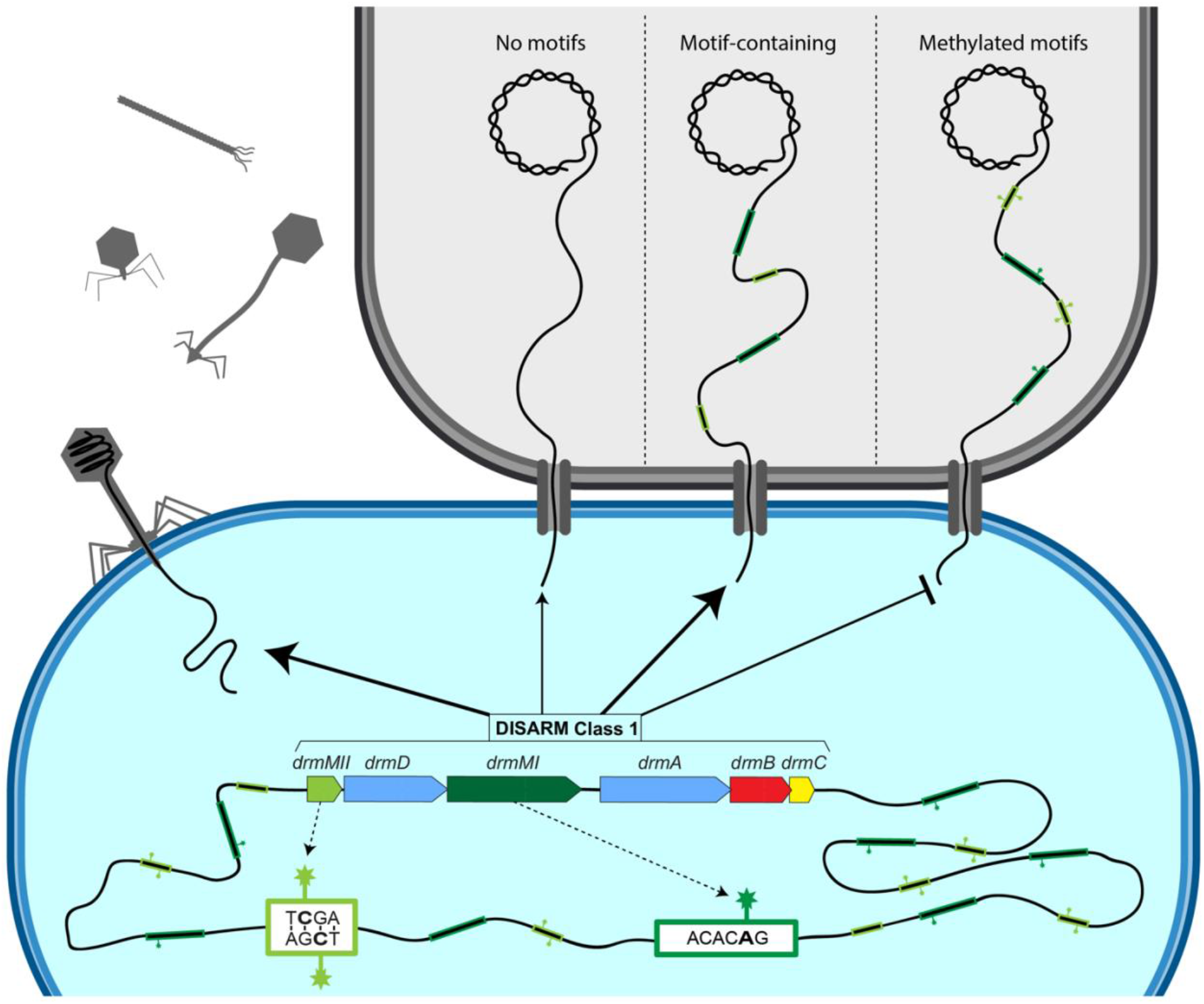
Mechanism of action of Class 1 DISARM of *Serratia sp.* SCBI. The methylases *drmMI* and *drmMII* methylate the host DNA at motifs ACAC<u>A</u>G and MT<u>C</u>GAK, respectively. The DISARM system provides protection from incoming unmethylated plasmid DNA and is less active on incoming DNA with methylated DISARM motifs. The efficiency of the protection increases with the number of unmethylated motifs present in the conjugated plasmid DNA.

## Supporting information

Aparicio-Maldonado_MS_BioRxiv.pdf

## ACKNOWLEDGEMENT

The authors thank Zohar Mukamel and the Weizmann Life Science Core Facilities for help with bisulfite and PacBio sequencing, respectively.

## AUTHOR CONTRIBUTIONS

S.J.J.B. and F.L.N. conceived the research. C.A.M. and A.S. performed the experiments. G.O. and R. S. performed the methylation-sensitive sequencing and corresponding data analysis. All authors contributed to data analysis and discussed the results. C.A.M. wrote the manuscript. F.L.N. and S. J.J.B. reviewed and edited the manuscript with input from all authors. All authors approved the manuscript.

