## Supplementary material for "Class I DISARM provides anti-phage and anti-conjugation activity by unmethylated DNA recognition": Aparicio-Maldonado_MS_BioRxiv.pdf

### Content

**Supplementary Table S1.** Bacteriophage features and host strains.

**Supplementary Table S2.** List of plasmids used in this work.

**Supplementary Table S3.** List of primers used in this work.

**Supplementary Table S4.** List of synthetic constructs used in this work.

**Supplementary Table S5.** Methylated sites of the strains determined by methylation-sensitive sequencing.

**Supplementary Table S6.** Methylation motifs found in phages and bacterial strains.

**Supplementary Figure S1.** Effect of phages on the growth curve of DISARM (+) or DISARM (-) strains.

**Supplementary Figure S2.** Growth of DISARM induced (DISARM +) and non-induced (DISARM -) strains compared to the wild-type strains.

**Supplementary Figure S3.** Effect of Class 1 DISARM on phage replication over time.

**Supplementary Figure S4.** Effect of Class 1 DISARM components on protection against T1 phage infection.

**Supplementary Figure S5.** Effect of Class 1 DISARM components on protection against phage infection.

**Supplementary Figure S6.** Conjugation efficiency of plasmid pCONJ with variable amount of DISARM motifs.

**Supplementary Figure S7.** Effect of DISARM on bacterial growth.

**Supplementary Table S1.** Bacteriophage features and host strains

| Phage | Replication strain | Infection type | Genetic material | Relevant genotype | Source |
|---|---|---|---|---|---|
| T1 | BL21 | Lytic cycle | dsDNA | ***dam+,*** *dcm-* | DSMZ |
| T3 | BL21 | Lytic cycle | dsDNA | *dam-*, *dcm-*, ***ocr+*** (demethylase) | DSMZ |
| T4 | BL21 | Lytic cycle | dsDNA | ***dam+***, *dcm-*, *hmc+*, *glc-hmc+* | Elizabeth Kutter's Lab |
| T7 | BL21 | Lytic cycle | dsDNA | ***ocr+*** (demethylase) | Ian Molineux's Lab |
| Lambda-vir | BL21 | Lytic cycle | dsDNA | *dam-*, *dcm-* | Elizabeth Kutter's Lab |
| Nami | Ci2 | Lytic cycle | dsDNA | *dam-*, *dcm-*, *hmc+*, | Brouns Lab collection |
| Shinka | Ci2 | Lytic cycle | dsDNA | ***dam+,*** *dcm-*, *hmc+*, | Brouns Lab collection |
| PhiX174 | C3000 | Lytic cycle | ssDNA | *dam-*, *dcm-* | DSMZ |
| M13 | JM109 | Chronic | ssDNA | *dam-*, *dcm-* | New England Biolabs |
| MS2 | WG5 | Lytic cycle | ssRNA | *dam-*, *dcm-* | DSMZ |

**Supplementary Table S2.** List of plasmids used in this work

| **Name in this study** | **Plasmid name** | **Insert** | **Backbone** | **Marker** | **Size (bp)** |
|---|---|---|---|---|---|
| pDIS_1 | pTU497 | *drmMII* | pET52_duet | Ampicillin | 6,165 |
| pDIS_2 | pTU496 | *drmD*, *drmMI* | pCOLA_duet | Kanamycin | 10,818 |
| pDIS_3 | pTU495 | *drmA*, *drmB*, *drmC* | pCDF_duet | Streptomycin | 10,276 |
| pDIS_4 | pTU501 | *drmMII*, *drmMI* | pCOLA_duet | Kanamycin | 8,777 |
| pDIS_5 | pTU498 | *drmA*, *drmB*, *drmC** | pCDF_duet | Streptomycin | 10,285 |
| pDIS_6 | pTU499 | *drmA*, *drmB**, *drmC* | pCDF_duet | Streptomycin | 10,285 |
| pDIS_7 | pTU500 | *drmA**, *drmB*, *drmC* | pCDF_duet | Streptomycin | 10,285 |
| pCONJ_0 | pTU503 | None | pSEVA331 | Chloramphenicol | 4,522 |
| pCONJ_1 | pTU504 | 2x ACACAG | pSEVA331 | Chloramphenicol | 4,517 |
| pCONJ_2 | pTU505 | 2x MTCGAK | pSEVA331 | Chloramphenicol | 4,522 |
| pCONJ_3 | pTU506 | 10x ACACAG | pSEVA331 | Chloramphenicol | 4,524 |
| pCONJ_4 | pTU507 | 10x MTCGAK | pSEVA331 | Chloramphenicol | 4,519 |
| pCONJ_5 | pTU508 | 2x ACACAG +2x MTCGAK | pSEVA331 | Chloramphenicol | 4,519 |
| pCONJ_6 | pTU511 | 10x ACACAG +10x MTCGAK | pSEVA331 | Chloramphenicol | 4,519 |

**Supplementary Table S3.** List of primers used in this work. Underlined the restriction site used for each primer. fw, forward primer; rv, reverse primer.

| **Name** | **Sequence (5'-3')** | **Description** | **fw/rv** |
|---|---|---|---|
| BN861 | TAAGGAGATATACCATGACTGATAACAACAAATCTAG | Clone *drmA,B,C* in pCDF-Duet | fw |
| BN1585 | CTGCGCTAGTAGACGAGCTTAGCCAAGAATCAAAACGATCGC | | rv |
| BN860 | GGTAAGGAGATATACCATGGAAATTTCTAATACACCTGATATCC | Clone *drmD,MI* in pCOLA-Duet | fw |
| BN2313 | GCCTAGGTTAATTAAGCTGGCAGCAGCCTAGGTTAATTAAGCTGCGCTAG | | rv |
| BN1219 | ACTTTAAGAAGGAGATATACCATGAAGAAAGTATCTTGTGTCG | Clone *drmMII* in pET-Duet | fw |
| BN2082 | GCAGCAGCCTAGGTTAATTAGTCATTCCACAGTGACCATATTCAC | | rv |
| BN1052 | ATTTTGTTTAACTTTAAGAAGGAGATATACCATGAAGAAAGTATCTTGTGTCG | Clone *drmMII* in pCOLA-Duet | fw |
| BN1053 | TAGTTATTGCTCAGCGGTGGCAGCAGCCTAGGTTAATTAGTCATTCCACAGTGACCATATTC | | rv |
| BN1056 | TATATATACATATGATGAGTATGGCCAGCATACACG | Clone *drmMI* in pCOLA-Duet | fw |
| BN1057 | ATATATATCTCGAGTTAGTTCGCACCATCAG[illegible]CAAATC | | rv |
| BN1539 | ACTTACATTCGCGAACTGCTG | Insert early stop codon in *drmA* of pTU495 to create pTU498 | fw |
| BN1540 | TTATCATTATTGATCCTCGATCTTTGGTATGTCTG | | rv |
| BN1542 | GCTGCTGTGCGAAAGGTATTAG | Insert early stop codon in *drmB* of pTU495 to create pTU499 | fw |
| BN1543 | TTATCATTAAAGAAGCCGGGCCTCTTGA | | rv |
| BN1545 | GAAAAGATACAAGCGATCGCTGGC | Insert early stop codon in *drmC* of pTU495 to create pTU500 | fw |
| BN1546 | TTATCATTAAGGAGAAACCAGACAAACTAGCTC | | rv |
| BN2208 | ATATATATGGATCCGTCTATCGGCATATGGTCAA | Amplify No-site module to clone with BamHI | fw |
| BN2209 | ATATATATGGATCCATATATCCTTTCGCCCGTCC | | rv |
| BN2218 | ATATATATGGTACCGTCTATCGGCATATGGTCAA | Amplify No-site module to clone with KpnI and NotI | fw |

| | | | |
|---|---|---|---|
| BN2393 | ATATATATGCGGCCGCATATATCCTTTCGCCCGTCC | | rv |
| BN2228 | ATATATATACTAGTCCGCATGGGCAATACGAGGT | Amplify 2 A-site module to clone with SpeI and BamHI | fw |
| BN2248 | ATATATATGGATCCAGCGGCATGGGTGACGGTTT | | rv |
| BN2230 | ATATATATACTAGTTAACCTAACGGTAAGAGGCT | Amplify 10 A-site module to clone with SpeI and BamHI | fw |
| BN2250 | ATATATATGGATCCTAAAGATTCATGTTAGTCCA | | rv |
| BN2228 | ATATATATACTAGTCCGCATGGGCAATACGAGGT | Amplify 2 A-site module to clone with SpeI and SphI | fw |
| BN2396 | ATATATATGCATGCAGCGGCATGGGTGACGGTTT | | rv |
| BN2230 | ATATATATACTAGTTAACCTAACGGTAAGAGGCT | Amplify 10 A-site module to clone with SpeI and SphI | fw |
| BN2397 | ATATATATGCATGCTAAAGATTCATGTTAGTCCA | | rv |
| BN2220 | ATATATATGGTACCTTTGAATACCTTATATTATT | Amplify 2 C-site module to clone with KpnI and NotI | fw |
| BN2394 | ATATATATGCGGCCGCCTGGTGACCGGTAGGTGTAC | | rv |
| BN2222 | ATATATATGGTACCGCGGATGGCGTTACAACTTT | Amplify 10 C-site module to clone with KpnI and NotI | fw |
| BN2395 | ATATATATGCGGCCGCACGGTCAGATTAGGCCCATG | | rv |
| BN2214 | ATATATATGGATCCTAACCGGCAATAATGCGC | Amplify pCONJ to insert modules using BamHI | rv |
| BN2215 | ATATATATGGATCCTACCAAATGCGGGACAAC | | fw |
| BN2226 | ATATATATGGTACCGAACATTGCGTTGCCTTG | Amplify pCONJ to insert modules using KpnI and NotI | rv |
| BN2390 | ATATATATGCGGCCGCAATGATGTCGTGGAACGTC | | fw |
| BN2234 | ATATATATACTAGTGAACATTGCGTTGCCTTG | Amplify pCONJ to insert modules using SpeI and SphI | rv |
| BN2391 | ATATATATGCATGCAATGATGTCGTGGAACGTC | | fw |

| | | | |
|---|---|---|---|
| BN2232 | ATATATATACTAGTTAACCGGCAATAATGCGC | Amplify pCONJ to insert modules using SpeI and BamHI | rv |
| BN2215 | ATATATATGGATCCTACCAAATGCGGGACAAC | | fw |
| BN2241 | TATATATATAAGGCGCGCCGAAGGCGTG | Amplify pSEVA331 backbone to clone Gblocks using AgeI and AscI | rv |
| BN2242 | TATATATATAACCGGTTATACCACCGTTGATATATCCCAATGGC | | fw |
| BN2243 | ATAATATCGGCGCGCCTC | Amplify Gblock_1 to clone using AscI and HindIII | fw |
| BN2244 | GACAGTCTGACAAGCTTCG | | rv |
| BN2245 | TATATATAGCGAAGCTTGGATAATTCC | Amplify Gblock_2 to clone using HindIII and AgeI | fw |
| BN2246 | TGAAAATAACCGGTGATTTTTTTCTC | | rv |

**Supplementary Table S4.** List of synthetic constructs used in this work, with cloning sites underlined and DISARM motifs in bold. Motifs in *italic* represent complementary sites for the non-palindromic site ACACAG.

| Name | Description | Sequence |
|---|---|---|
| Gblock_1 | Sequence modification to replace putative DISARM motifs in backbone sequence. Cloned using AscI and HindIII | ATAATATCGGCGCGCCTCCCCCTCCGGCAAAAAGTGGCCC<br>CTCCGGGGCTTGTTGATGGACTGCGCGGCCTTCGGCCTTGC<br>CCAAGGTGGCGCTGCCCCCTTGGAACCCCCGCACTCGCCG<br>CCGTGAGGCTCGGGGGGCAGGCGGGCGGGCTTCGCCCTTG<br>GACTGCCCCCACTCGCATAGGCTTGGGTCGTTCCAGGCGC<br>GTCAAGGCCAAGCCGCTGCGCGGTCGCTGCGCGAGCCTTG<br>ACCCGCCTTCCACTTGGTGTCCAACCGGCAAGCGAAGCGC<br>GCAGGCCGCAGGCCGGAGGCTTTTCCCCAGAGAAAATTAA<br>AAAAATTGATGGGGCAAGGCCGCAGGCCGCGCAGTTGGAG<br>CCGGTGGGTATGTGGTAGAAGGCAGGGTAGCCGGTGGGCA<br>ATCCCAGTGGTCAAGCTCGTGGGCAGGCGCAGCCTGTCCA<br>TCAGCTTGTCCAGCAGGGTTGTCCACGGGCCGAGCGAAGC<br>GAGCCAGCCGGTGGCCGCTCGCGGCCATCGTCCACATATC<br>CACGGGCTGGCAAGGGAGCGCCGCGACCGCGCCGGGCGA<br>AGCTTGTCAGACTGTC |
| Gblock_2 | Sequence to insert the YFP and MAM into the backbone sequence. Cloned using HindIII and AgeI | TATATATAGCGAAGCTTGGATAATTCCTAATTTTTGTTGAC<br>ACTCTATCGTTGATAGAGTTATTTTACCACTCCCTATCAGT<br>GATAGAGAAAAGAATTCAAAAGATCTAGGAGGAAAAAAA<br>TGGTGAGCAAGGGCGAGGAGTTGTTCACCGGGGTGGTGCC<br>CATCCTGGTTGAGCTGGACGGCGACGTAAACGGCCACAAA<br>TTTTCCGTGTCCGGCGAGGGCGAGGGCGATGCCACCTACG<br>GCAAGCTGACCCTGAAGCTGATCTGCACCACCGGCAAGCT<br>GCCCGTGCCCTGGCCCACCCTCGTGACCACCTTGGGCTACG<br>GCCTGCAGTGCTTCGCCCGCTACCCCGACCACATGAAGCA<br>ACACGACTTCTTCAAGTCCGCCATGCCCGAAGGCTACGTCC<br>AGGAGCGCACCATCTTCTTCAAGGACGACGGCAACTACAA<br>GACCCGCGCCGAGGTGAAGTTTGAGGGCGACACCCTGGTG<br>AACCGCATTGAACTGAAGGGCATTGACTTCAAGGAGGACG<br>GCAACATCTTGGGGGCCCAAGCTGGAAGTACAACTACAAC<br>CGCCACCACGTCCATATCACCGCCGACAAGCCGAAGAACG<br>GCATCAAGGCCAACTTCAAGATCCGCCACCACCTCAAGGA<br>CGGCGGCGTGCAGCTCGCCGACCACTACCAGCAGAACACC<br>CCCATCGGCGACGGCCCCGTGCTGCTGCCCGACAACCACT<br>ACCTAAGTAAGGATCTCCAGGCATCAAATAAAACGAAAGG<br>CTCAGTTGAAAGACTGGGCCTTTCGTTTTATGAGCAAGCCC |

| | | |
|---|---|---|
| | | GTAGGGGCCAGTCTAGCTTCAAGTATGACGGGCTGATACT<br>GGGCCGGCAGGCGCTCCATTGCCCAGTCGGCAGCGACATC<br>CTTCGGGCGCATTATTGCCGGTTACTGCGCTGTACCAAATG<br>CGGGACAACGTAAGCACTACATTTCGCTCATCGCCAGCCC<br>AGTATCTGATGCACGGGCGGCGAGTTCCATAGCGTTAAGG<br>TTTCATTTAGCGCCTCAAATAGATCCTGTTCAGGAACCGGA<br>TCAAAGAGTTCCTCCGCCGCTGGACCTACCAAGGCAACGC<br>AATGTTCTCACGGAATGATGTCGTGGAACGTCAACAATGG<br>TGACTTCTACAGCGCGGAGAATCTCGCTCTCTCCAGGGGTT<br>GCTTTTGTCAGCAAGAGCCCAGCAGTTTAATTAAAGCGGA<br>TAACAATTTCTCTCGGGAGGCCGCCTAGGCCCGTCGTGACT<br>GGGAAAACCCTGGCGACGTGGGTCCCCAATAATTACGATT<br>TAAATTGGCGAAAATGAGACGTTGATCGGCACGTAAGAGG<br>TTCCAACTTTCACCATAATGAAATAAGATCACTACCGGGCG<br>TATTTTTTGAGTTATGGAGATTTTCAGGAGCTAAGGAAGCT<br>AAAATGGAGAAAAAAATCACCGGTTATTTTCA |
| Gblock_3 | Module containing No motifs (for control) | GTCTATCGGCATATGGTCAATTTGAGTGTCCGGAGGCGGA<br>AATCCGCCACGAATGAGAATGTATTTCCCCGACAATCATA<br>ATGGGGCGCTCCTAAGCTTTTCCATTGGTTGGGCCGGCTAG<br>GCCATCATCTGCCCGGAGTTTCGGCGCACTGCTGCCGACAT<br>GCCGGGCATTGTTTTAGGGGCGTTATTCTTGAGGGCACTCG<br>GAGCTAATTTGTCGCGACCAGCCGGGGTAGTCATCGGGCT<br>TATACACGAAAAGCCCATGTATCCGTTTCCACGTATGGAAC<br>GTCTTTAGCTCCGGCAAGCAATTAAGAACAACGCAAGCGG<br>GACGGGCGAAAGGATATAT |
| Gblock_4 | Module containing 2 C motifs | TTTGAATACCTTATATTATTCATCGCGGATATAAACATGAG<br>AAACGGCCGAATACACCATGTTCGTATCGTATCGGTAAAT<br>ACCTCGCGTAGCCATGTGCCATATGGTTGCGA**TCGA**TTGGT<br>TATGCATATGGTCCACATGGACACTCTTCGCTTCCGGGTAT<br>GCGCTATATGTGACGGTCTTTAGGCGCACTGATGCTCATGC<br>**TCGA**GTTAAACCAACCGACACCAGATCATGTAAGGTCCGC<br>CACGCATGACGACATGCCCACGGAGATCACCGACCGATCT<br>ATCTGATCGGCGACCATTTGTGTGTTACTGGGGCGGAGGCT<br>GGTGACCGGTAGGTGTAC |
| Gblock_5 | Module containing 10 C motifs | GCGGATGGCGTTACAACTTTCCTTAATGGCTCTTGGGCCGC<br>GGTGCGTGACCTTGCAGGAATTGAC**TCGA**GCCGTTAATTT<br>CCCTTGCATACAA**TCGA**TGTTTTTTTGTCTTTTTATCCGCTT<br>C**TCGA**GATAAGAGTGACATACTTCTTAA**TCGA**TCGCCTCC<br>GTACACATGTACGATCGC**TCGA**GCCATGAGAATAGGTATA<br>CCATGTA**TCGA**TGTGAGCAACGAAAGCCTAAACGGGAC**TC** |

| | | |
|---|---|---|
| | | **GA**GCGGCCAAAAGTCGGTGCGAAT**TCGA**GTCA**TCGA**TATGTTGGTCTGGCTATGATCTACC**TCGA**GCAGGCGTTACGACGGTCAGATTAGGCCCATG |
| Gblock_6 | Module containing 2 A motifs | CCGCATGGGCAATACGAGGTCGTCTGCTCTGGTCAGCCTCTAATGGCTCGTTAGATAGTCTAGCCGCTGGTAATCACATGACCTCGTCTCCCCATTGGTGCTACGGCGATTCTTGGAG**ACACAG**GCTGCGATCGCTAATGTGAGGACATGTGTAATATTAGC***CTGTGT***TAAGTCCCCAATGGTTGTGGCCTTTTGAAAAGTGAACTTCATAACATATGCATGTCTCACGCACATGGATGGTTTGGACAAATTTGATTCAAGTCTGATCAACCTTCACACATGATCTAGAATCAAAAGCATGATCTCCCGGGTGCGAAATAAGCGGCATGGGTGACGGTTT |
| Gblock_7 | Module containing 10 A motifs | TAACCTAACGGTAAGAGGCTCAAAATACTACGTAAC**ACACAG**GACTGCGACGTTCTAAA***CTGTGT***TCCGTCTGAACCGCCATCCATGGATCAC**ACACAG**CCGAAAAAAAGATATCAGGAACT***CTGTGT***CTCACATCGGTCATATGGAAACTACATGG**ACACAG**CTTCCTGGCAACCGGGGGGTGGTAATCCG***CTGTGT***ATGAGAAGGTATTTGCTCAATAATCA**ACACAG**CAGGCATCTAACTTTTCCCATGCC***CTGTGT***CGGCTTGCCCATTTTGCCATGTA**ACACAG**TTAGGACTCGCGCCAACGCGCA***CTGTGT***ATTCGGTAAAGATTCATGTTAGTCCA |

**Supplementary Table S5.** Methylated sites of the strains determined by methylation-sensitive sequencing. m6A refers to ACACAG sites and 5mC to MTCGAK sites.

| **Strain** | **DISARM genes** | **Number of m6A sites detected** | **Ratio of m6A sites (%)** | **Number of 5mC sites detected** | **Ratio of 5mC sites (%)** | **Number of GmATC sites detected** | **Ratio of GmATC sites (%)** |
|---|---|---|---|---|---|---|---|
| WT | - | 0 | 0 | 0 | 0 | 37377 | 99.9 |
| DISARM | *drmMII, drmD, drmMI, drmA, drmB, drmC* | 929 | 84.3 | 1201 | 67 | 37359 | 99.9 |
| DrmMII-DrmMI | *drmMII, drmMI* | 815 | 73.9 | 794 | 36 | 37374 | 99.9 |
| DrmMII | *drmMII* | 0 | 0 | 234 | 34 | 37375 | 99.9 |
| DrmMI | *drmMI* | 619 | 56.1 | 0 | 0 | 37375 | 99.9 |

**Supplementary Table S6:** Methylation motifs found in phages and bacterial strains

| Phage/ bacterial strain | Genome size (bp) | Number of ACACAG sites | % ACACAG sites per 100 kb | Expected ACACAG sites | Number of MTCGAK sites | % MTCGAK sites per 100 kb | Expected MTCGAK sites | Number of TCGA sites | % TCGA sites per 100 kb | Expected TCGA sites |
|---|---|---|---|---|---|---|---|---|---|---|
| T1 | 48,836 | 12 | 147 | 12 | 10 | 164 | 12 | 202 | 1,655 | 191 |
| T3 | 38,209 | 12 | 188 | 9 | 8 | 167 | 9 | 134 | 1,403 | 149 |
| T4 | 168,903 | 36 | 128 | 41 | 24 | 114 | 41 | 312 | 739 | 660 |
| T7 | 39,937 | 18 | 270 | 10 | 8 | 160 | 10 | 111 | 1,112 | 156 |
| Lambda-vir | 48,502 | 19 | 214 | 12 | 10 | 165 | 12 | 121 | 998 | 189 |
| Nami | 165,679 | 33 | 120 | 40 | 26 | 126 | 40 | 311 | 751 | 647 |
| Shinka | 162,160 | 37 | 137 | 40 | 21 | 104 | 40 | 281 | 693 | 633 |
| M13 | 7222 | 1 | 83 | 2 | 2 | 222 | 2 | 11 | 1,218 | 28 |
| PhiX174 | 5,386 | 1 (rev[a]) | 111 | 1 | 1 | 149 | 1 | 10 | 743 | 21 |
| *Serratia* sp. SCBI | 5,034,688 | 717 | 85 | 1,229 | 1,399 | 222 | 1,229 | 24,114 | 1,916 | 19,667 |
| *Serratia* plasmid | 67,208 | 19 | 170 | 16 | 37 | 440 | 16 | 263 | 1,565 | 263 |
| *Serratia* ATCC 39006 | 4,971,757 | 1783 | 221 | 1,214 | 946 | 152 | 1,214 | 13,376 | 1,076 | 19,421 |
| *E. coli* Dh5α | 4,583,637 | 1,132 | 148 | 1,119 | 1001 | 175 | 1,119 | 15,292 | 1,334 | 17,905 |
| *E. coli* BL21-AI | 4,530,564 | 1,106 | 146 | 1,106 | 993 | 176 | 1,106 | 14,937 | 1,319 | 17,698 |
| *E. coli* K12 JM109 | 4,641,652 | 1150 | 149 | 1,133 | 1,013 | 175 | 1,133 | 15,462 | 1,332 | 18,131 |
| *E. coli* JM109(DE3) | 4,661,885 | 1159 | 149 | 1,138 | 108 | 19 | 1,138 | 15,409 | 1,322 | 18,210 |
| *E. coli* C3000 | 4,617,024 | 1,134 | 147 | 1,127 | 1,010 | 175 | 1,127 | 15,575 | 1,345 | 18,035 |
| *E. coli* WG5 | 4,592,887 | 1,127 | 147 | 1,121 | 1,002 | 175 | 1,121 | 15,449 | 1,345 | 17,941 |

[a] rev: site present in complementary strand of the ssDNA phage genome.

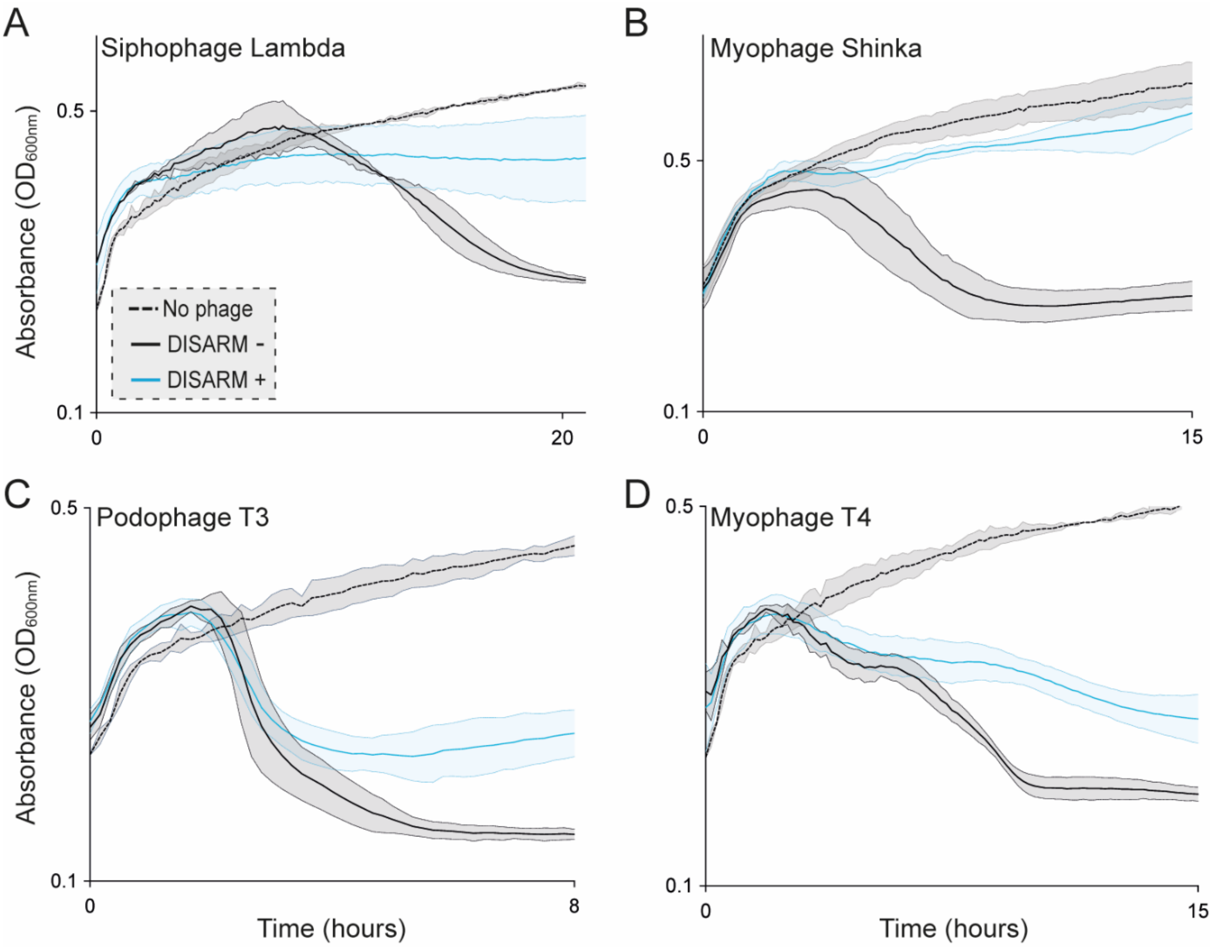


**Supplementary Figure S1.** Effect of phages on the growth curve of DISARM (+) or DISARM (-) strains. (**A**) Siphophage Lambda, (**B**) Myophage Shinka, (**C**) Podophage T3, and (**D**) Myophage T4. Lines and filled areas indicate average and standard deviation of three independent replicates, respectively.

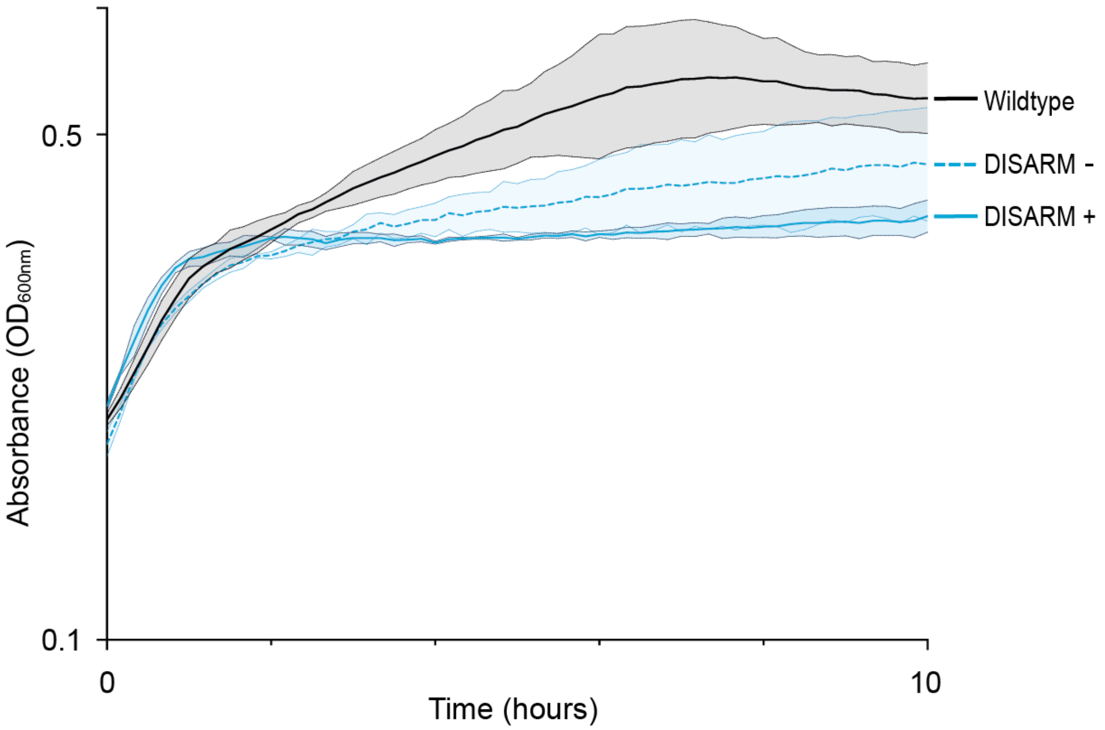


**Supplementary Figure S2.** Growth of DISARM induced (DISARM +) and non-induced (DISARM -) strains compared to the wild-type strains. Line and filled areas within dotted lines indicate average and standard deviations of three independent replicates, respectively.

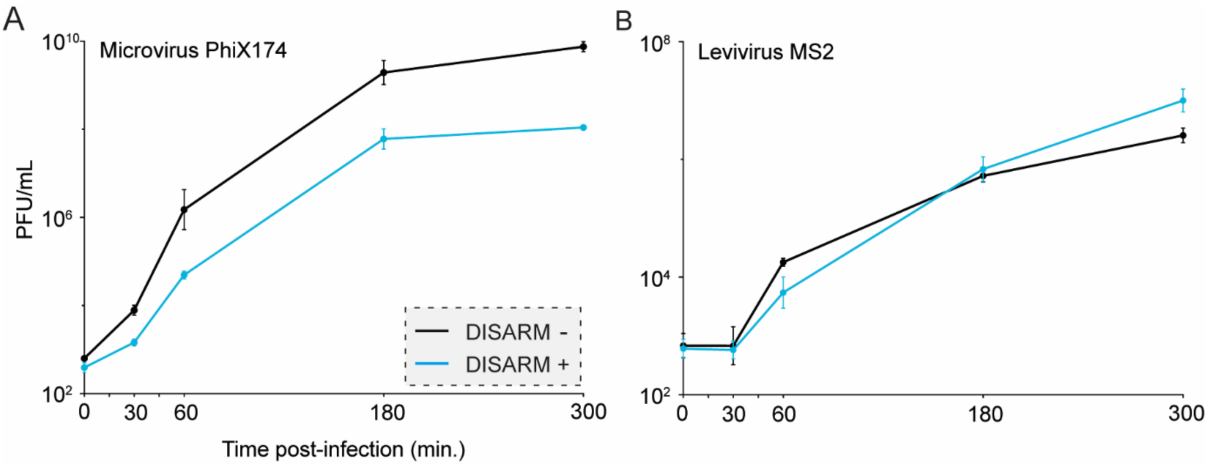


**Supplementary Figure S3.** Effect of Class 1 DISARM on phage replication over time. (**A**) Phage phiX174. (**B**) Phage MS2. Bacterial cultures of DISARM (+) or DISARM (-) strains were infected with phage at an MOI of $10^{-6}$ and the titer determined at selected time points.

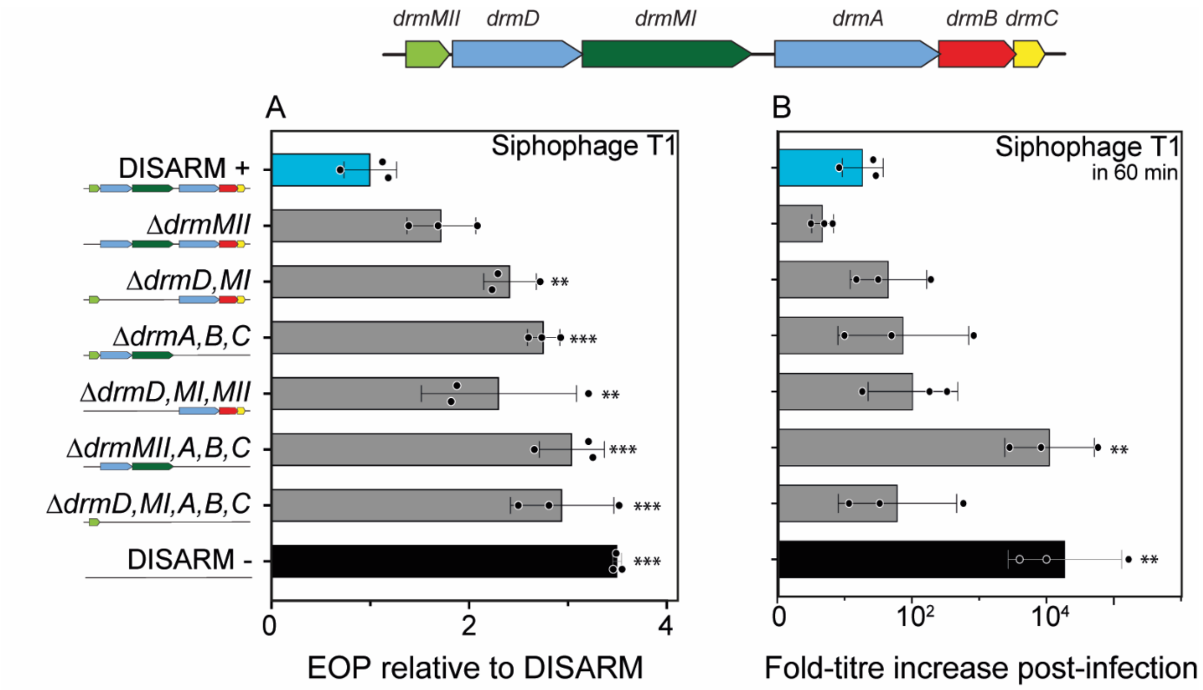


**Supplementary Figure S4.** Effect of Class 1 DISARM components on protection against T1 phage infection. (**A**) Efficiency of plating (EOP) of siphophage T1 on strains containing the modified DISARM system (DISARM (-) strain), normalized to the DISARM strain. (**B**) Titer-fold increase of siphophage T1 upon propagation in cultures of strains containing the complete or modified DISARM system. Graphic represents the time point at which maximum effect on phage replication was achieved. Bars depict the average and standard deviation of three independent replicates. Statistical significance was determined by one-way ANOVA+ with Tukey post-hoc test and is represented as *, **, or *** for $p < 0.05$, 0.01 or 0.001, respectively.

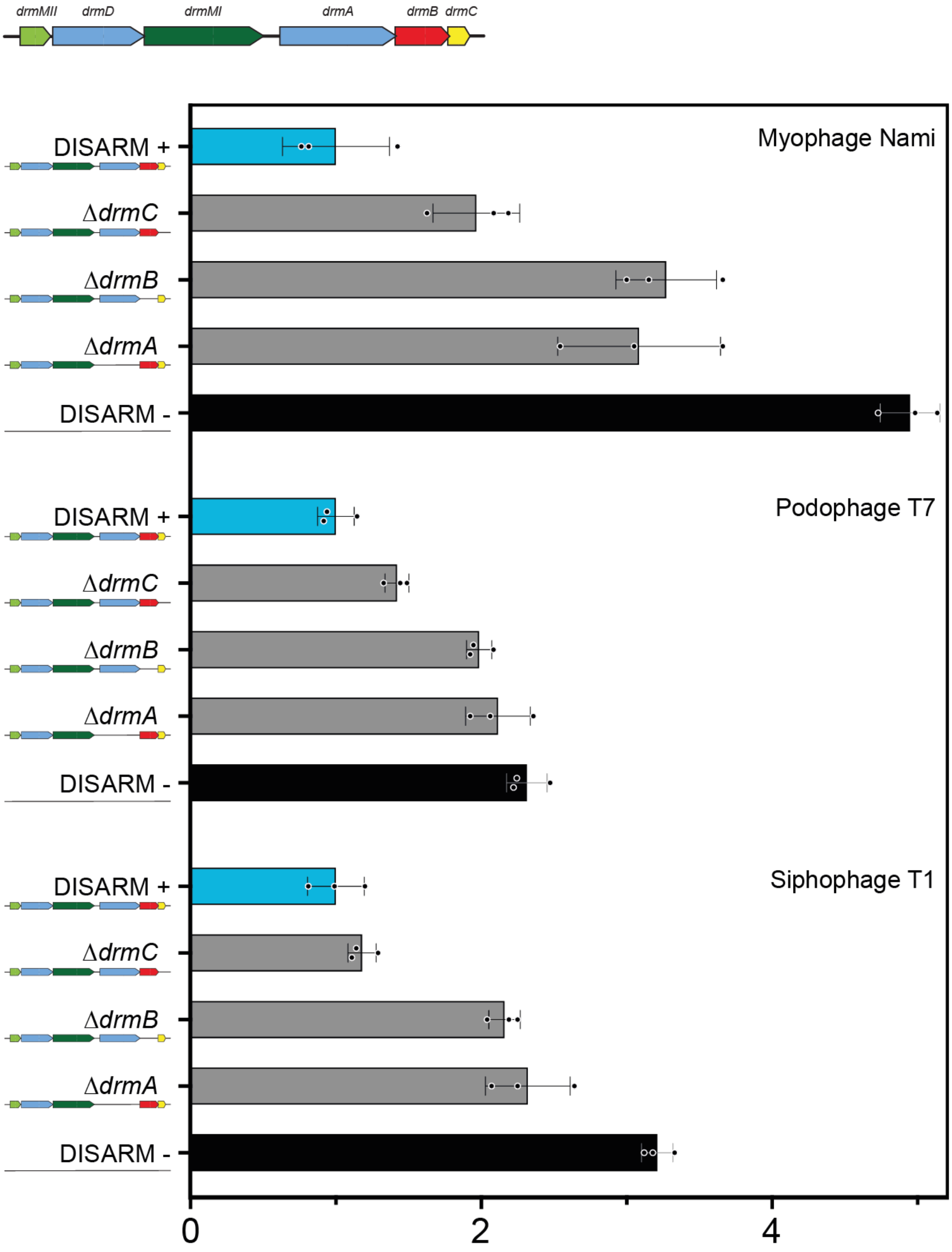


**Supplementary Figure S5.** Effect of Class 1 DISARM components on protection against phage infection. Efficiency of plating (EOP) of myophage Nami (top), podophage T7 (mid), and siphophage T1 (bottom) on strains containing the DISARM system with deletions of genes *drmA*, *drmB*, *drmC*; normalized to the DISARM (+) strain.

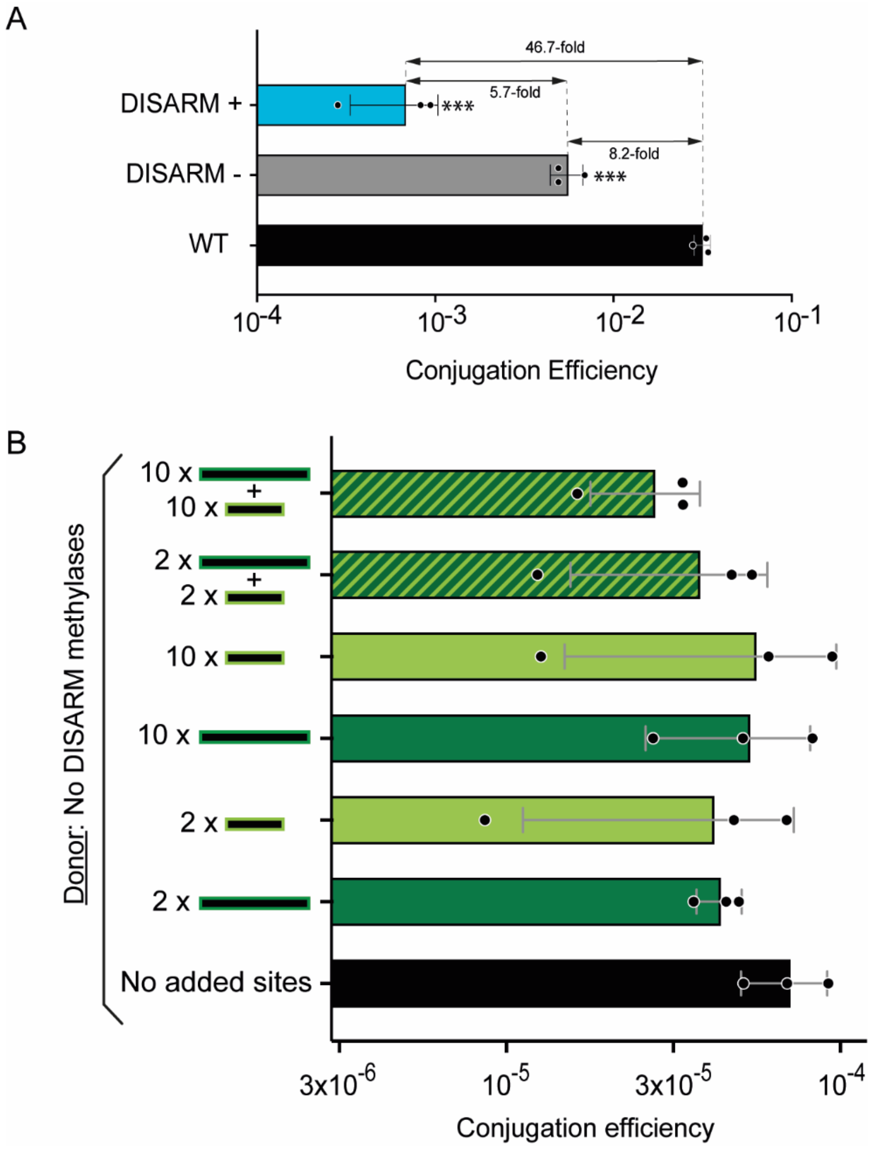


**Supplementary Figure S6.** Conjugation efficiency of plasmid pCONJ with variable amount of DISARM motifs. (**A**) Conjugation efficiency of plasmid pCONJ without motifs in DISARM induced (DISARM+), non-induced (DISARM -) and control strains. (**B**) Conjugation efficiency of plasmid pCONJ with variable amounts of unmethylated motifs ACACAG (dark green) and/or MTCGAK (light green) into the recipient control strain.

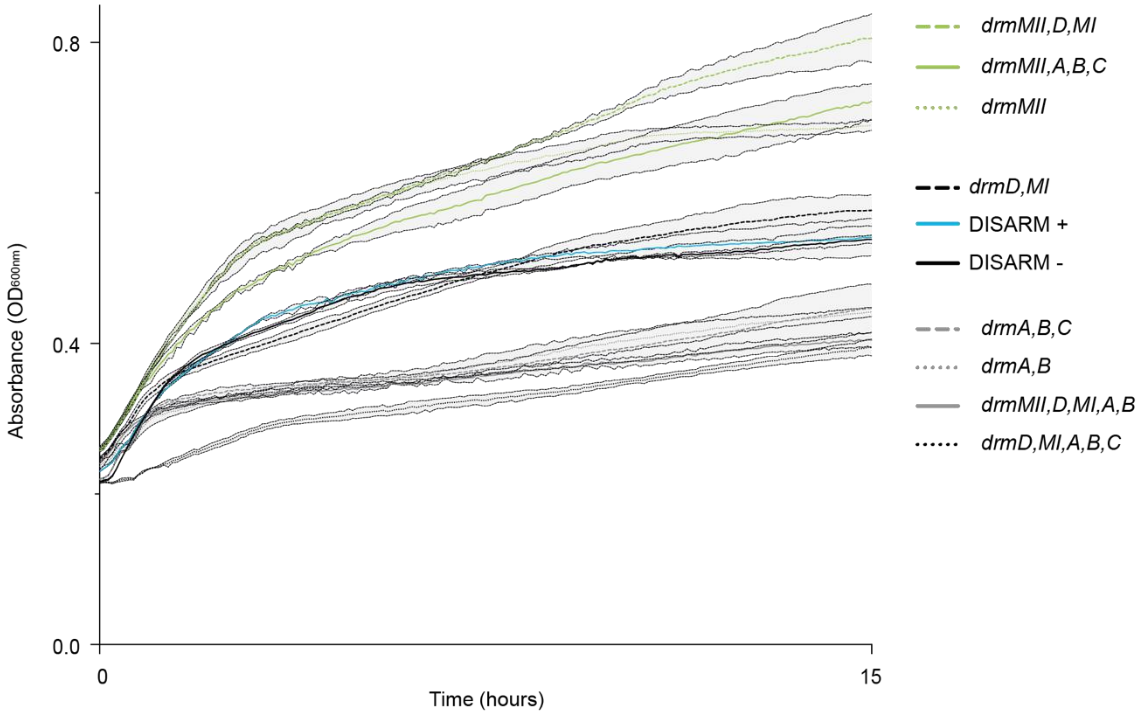


**Supplementary Figure S7.** Effect of DISARM on bacterial growth. Lines and filled areas within dotted lines indicate average and standard deviations of three independent replicates, respectively.
